# Computed tomography lacks sensitivity to image gold labelled mesenchymal stromal cells in vivo as evidenced by multispectral optoacoustic tomography

**DOI:** 10.1101/2022.06.15.495483

**Authors:** Alejandra Hernandez Pichardo, James Littlewood, Arthur Taylor, Bettina Wilm, Raphaël Lévy, Patricia Murray

**Author notes:** Correspondence should be addressed to Patricia Murray.

## Abstract

Elucidating the mechanisms of action and long-term safety of cell therapies is necessary for their clinical translation. Non-invasive imaging technologies such as bioluminescence imaging (BLI), computed tomography (CT) and multispectral optoacoustic tomography (MSOT) have been proposed as tools for longitudinal cell monitoring but their performances have not been compared. Here, we evaluate combinations of these modalities to track the in vivo distribution of gold-labelled mesenchymal stromal cells (MSCs). We found that injected MSCs labelled with gold nanoparticles and expressing the reporter gene firefly luciferase could be detected with BLI and MSOT but not CT. We conclude that the MSCs did not carry enough contrast agent to be tracked by CT, demonstrating that CT tracking of gold-labelled cells is not a practical approach as high amounts of gold, which might impair cell viability, are necessary.

## 1. Introduction

The ability to image cells and non-invasively track their fate in animal models has become increasingly important in assessing the long-term safety and efficacy of cell-based regenerative therapies. Moreover, the ability to monitor the biodistribution of cells over time can also offer key insights into their mechanisms of action; for instance, establishing whether engraftment in the target organ is required for the cells to have any beneficial effects.

The intravenous administration of mesenchymal stromal cells (MSCs) in mice leads to their entrapment in the pulmonary vasculature and inability to reach other organs (1–3). Bioluminescence imaging (BLI) has been key in this discovery. BLI is a non-invasive, whole-animal, pre-clinical imaging modality with high sensitivity and a temporal resolution of seconds to minutes (4). BLI allows longitudinal cell tracking via a reporter gene encoding luciferase, an enzyme that oxidises a substrate to generate light (5,6). However, BLI is limited by low spatial resolution (3 – 5 mm) which does not allow the biodistribution of the cells to be mapped at the intra-organ level (7).

Multispectral optoacoustic tomography (MSOT) is a non-invasive imaging modality that provides functional and anatomical information in real-time. MSOT operates by the photoacoustic effect: incident modulated light energy is absorbed leading to thermo-elastic expansion and the generation of ultrasound waves (8). MSOT uses a range of near-infrared excitation wavelengths, and subsequent spectral unmixing algorithms allows the identification of the optical signatures of endogenous and administered contrast agents. It benefits from high spatial (100 µm) and temporal (0.1 s) resolutions. Its main limitation for tracking cells delivered intravenously is that due to the high air content of the lungs and the behaviour of sound in this medium, MSOT is unable to image this organ.

Computed tomography (CT) is inherently effective for lung imaging due to the native contrast provided by the airspaces. CT is a non-invasive imaging modality that generates 3D anatomical images based on the differential X-Ray attenuation of materials (9). Its high spatial resolution (50-200 µm) with high signal-to-noise ratio, high depth of penetration, quantitative capabilities, fast temporal resolution, and cost-effectiveness have made it a widely applied imaging modality in the clinic (10). The main drawbacks of CT are that it uses ionising radiation, and its sensitivity is low (10,11).

To distinguish administered cells from endogenous tissue in animal models, cells need to be labelled with a contrast agent. Gold nanoparticles, which exist in different shapes, are a suitable contrast agent for both CT and MSOT. The CT contrast is due to gold’s high density and atomic number (9) whilst MSOT contrast can be achieved via the near-infrared (NIR) longitudinal surface plasmon bands that are characteristic of gold nanorods (GNRs) (12). Cell labelling with GNRs is achieved by endocytosis. The tight packing of the GNRs within endosomes can result in the GNRs undergoing plasmon coupling, altering their optical properties and compromising MSOT detection (13). Coating the GNRs with silica prevents plasmon coupling and does not affect cell viability, allowing MSOT to reach its full potential (14).

Our group has previously applied a dual BLI/MSOT imaging strategy to track GNR-labelled cells (13,15). Here, the possibility of expanding this approach to include CT is explored. In addition, we aimed to compare the effectiveness of MSOT and CT for tracking cells labelled with silica-coated GNRs to study the in vivo biodistribution of MSCs delivered subcutaneously or intravenously.

## 2. Materials and Methods

### 2.1 Nanoparticle characterization

Commercially available silica-coated gold nanorods (GNRs) pre-adsorbed with bovine serum albumin (BSA) were purchased from Creative Diagnostics (2.5 mg/mL). Their properties were characterized using transmission electron microscopy (TEM) (Tecnai G2 Spirit BioTWIN) coupled to a Gatan RIO16 camera and Vis-NIR spectroscopy (FLUOstar Omega, BMG Labtech). GNR stability was studied by incubating the GNRs in cell culture medium at 37°C in a humidified incubator, with 5% CO_2_. After 24 h, the GNRs were recovered by centrifugation at 13000 x g for 20 min, washed three times with dH_2_O and imaged by TEM. Particles were deposited onto glow discharged fomvar/carbon coated grids for 10 mins, excess wicked off and stained for 20 s with 1% aqueous uranyl acetate.

### 2.2 Cell isolation, generation of reporter cell line and culture

Human umbilical cord-derived mesenchymal stromal cells (hUC-MSCs) were obtained from the National Health Service Blood and Transplant (NHSBT, UK) at passage 3 (p3). The hUC-MSCs were transduced with a lentiviral vector encoding luc2 firefly luciferase (FLuc) reporter under the constitutive elongation factor 1-α (EF1α) promoter and the ZsGreen fluorescent protein downstream of the bioluminescence reporter via an IRES linker. The pHIV-Luc2-ZsGreen vector was kindly gifted by Bryan Welm and Zena Werb (Addgene plasmid #39,196) (16). To obtain a >98% FLuc positive population, the cells were sorted based on ZsGreen fluorescence.

The cells were grown in MEM-α and supplemented with 10% foetal bovine serum (FBS) (Gibco) and kept at 37°C in a humidified incubator, with 5% CO_2_.

### 2.3 Cell viability assay

5 × 10^3^ cells were seeded into 96-well plates (Corning) and allowed to attach for 24 h. The viability of hUC-MSCs after 24 h exposure to increasing concentrations of GNRs was determined by the CellTiter-Glo™ Luminescent Cell Viability Assay (Promega Corporation), which generates luminescent signals based on ATP levels. Tests were performed in triplicate with two PBS washing steps between GNR exposure and the assay. Luminescence was measured in a multi-well plate reader (FLUOstar Omega, BMG Labtech).

### 2.4 Assessing the extent of GNR uptake by hUC-MSCs

hUC-MSCs were seeded at 13 × 10^3^ cells/cm^2^ into 24-well plates (Corning) and allowed to attach for 24 h. Based on the available material, cells were exposed to 1:100 and 1:10 GNR dilutions (0.125 or 0.25 mg/mL) in cell culture medium for 24 h. After this period, the cells were washed with PBS to remove excess GNRs and fixed with paraformaldehyde (4 % w/v in PBS, pH 7) for 20 min at room temperature (RT). GNR uptake by the cells was assessed by using a silver enhancement solution kit (Sigma SE100) according to the manufacturer’s instructions. After rinsing three times with PBS, the cells were imaged by light microscopy with a Leica DM IL microscope coupled to a DFC420C camera.

### 2.5 Animal experiments

Eight-to ten-week-old female albino (C57BL/6) (B6N-TyrC-Brd/BrdCrCrl, originally received from the Jackson Lab) mice were used for all animal experiments. Mice were housed in individually ventilated cages (IVCs) under a 12-h light/dark cycle and provided with standard food and water ad libitum. All animal procedures were performed under a licence granted under the Animals (Scientific Procedures) Act 1986 and were approved by the University of Liverpool Animal Welfare and Ethics Review Board (AWERB).

Mice were injected with 5 × 10^5^ FLuc-hUC-MSCs (hUC-MSCs hereinafter) suspended in 100 μL of PBS by either intravenous (IV) or subcutaneous (SC) administration to the flank, and subsequently imaged via BLI, MSOT and CT, all under terminal anaesthesia with isoflurane.

### 2.6 Bioluminescence imaging

Immediately after cell injection, the animals received a SC injection of D-Luciferin (10 μL/g [body weight] of a 47 mM stock solution). 20 min later, the animals were imaged with an IVIS Spectrum instrument (Perkin Elmer). All data is displayed in radiance (photons/second/centimeter^2^/steradian), where the signal intensity scale is normalised to the acquisition conditions.

### 2.7 Multispectral optoacoustic tomography imaging

All imaging was performed in the inVision 256-TF MSOT imaging system (iThera Medical, Munich, Germany).

Tissue-mimicking imaging phantoms with a 2 cm diameter were constructed from 1.5 w/v % agar and 0.4 w/v % intralipid in distilled water (17). Two cavities were created to facilitate insertion of clear straws containing either unlabelled hUC-MSCs or hUC-MSCs labelled with 0.25 mg/mL GNRs. 5 × 10^5^ hUC-MSCs were prepared by labelling and trypsinization as described above and suspended in 100 μL of PBS. Then, the whole volume was inserted into the phantom cavity.

The agar phantoms with inserts were imaged at 61 wavelengths (680 nm to 980 nm in 5 nm steps) at 25°C. 3 frames were measured per wavelength and averaged.

To image mice in vivo, their abdomens were shaved and de-epilated using Veet Hair removal cream (Reckitt Benckiser, UK) 24 h before imaging. Mice were imaged at 34°C. In mice receiving subcutaneous hUC-MSC injection, scans were acquired at the site of injection at 61 wavelengths (680 nm to 980 nm in 5 nm steps) in 1mm slices. 10 frames per wavelength were measured and averaged. In mice receiving intravenous hUC-MSC injection, scans were acquired at the lungs at 61 wavelengths (680 nm to 980 nm in 5nm steps) in 1 mm slices. Additionally, images were acquired through the full volume of all animals at 8 wavelengths (660, 700, 730, 750, 760, 800, 850, 900 nm) in 1 mm slices.

For image processing, the ViewMSOT 4.0.1.34 (iThera Medical, Germany) was used. Data were reconstructed with the back-projection algorithm. Multispectral unmixing was performed using the linear regression algorithm. Images were unmixed for haemoglobin, oxyhaemoglobin, melanin, and the GNR MSOT spectrum.

### 2.8 Computed Tomography

Agar phantoms with inserts, as used for MSOT, were imaged using an aluminium filter 0.5 mm thick or a 0.06 mm copper filter with an applied X-ray tube voltage of 90 kV in a Quantum GX micro CT (Rigaku Corporation). Images were acquired with a field of view (FOV) of 25 mm giving a voxel size of 50 µm.

After MSOT imaging, the mice were culled and their carcasses were imaged using an aluminium filter 0.5 mm thick and an applied X-ray tube voltage of 90 kV with the same instrument. Images were acquired with a FOV of 25 mm giving a voxel size of 50 µm. Surface-rendered 3D models were constructed for 3D viewing of the analysed mice. Volume rendered 3D images were generated using the Quantum GX software version 3.0.39.5100.

### 2.9 Statistical analysis

All values in graphs are represented as mean ± standard deviation. The statistical analysis was performed using the GraphPad Prism software. The type of statistical test and the number of replicates included in the analyses are indicated in the figure legends.

## 3 Results

### 3.1 Gold nanorod characterisation

To characterise the silica-coated GNRs, their absorbance spectrum was assessed using Vis-NIR spectroscopy which revealed that their longitudinal surface plasmon resonance (LSPR) peaks at 738 nm. The integrity of the silica shell was evaluated after incubation in cell medium as etching might occur (18). After 24 h, the GNRs lose the silica coating resulting in a 25 nm LSPR left shift with a peak at 713 nm (Figure 1a).

**Figure 1.**
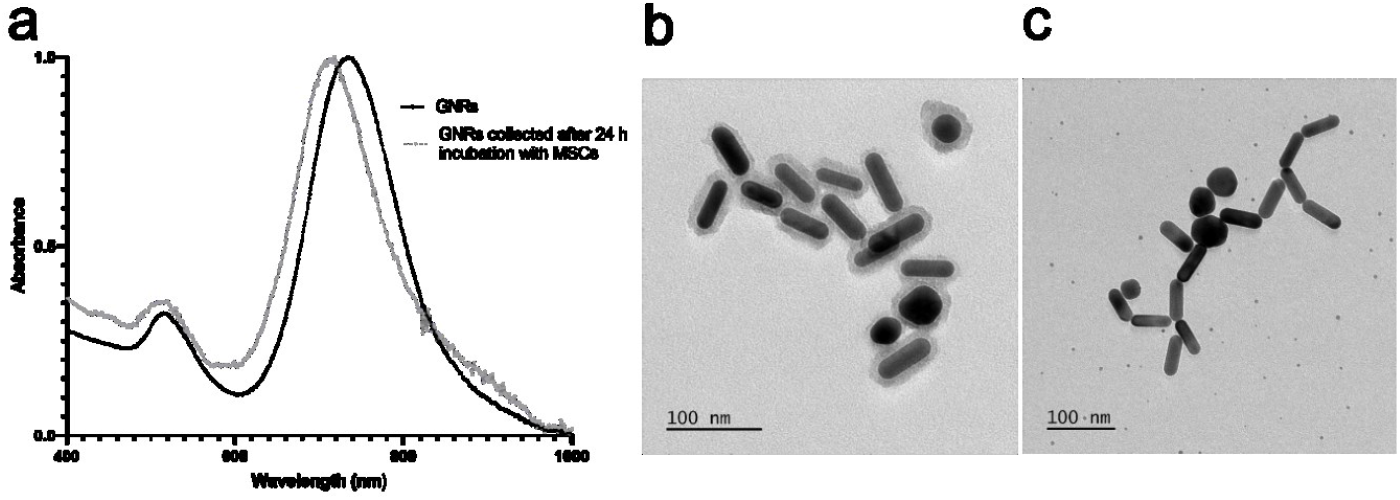
Characterisation of GNRs. (a) Vis-NIR spectrum of GNRs. (b) Representative TEM picture of silica-coated GNRs. (c) TEM image of GNRs 24 h post incubation in cell culture medium.

The size of the GNRs and the thickness of the silica shell was determined using transmission electron microscopy (TEM). The core size was 55.77 ± 7.32 nm length by 17.36 ± 1.99 nm width with a silica shell thickness of 7.25 ± 1.65 nm. TEM confirmed the loss of the silica shell after incubation in cell medium (Figure 1b). Despite the LSPR shift, the GNRs absorbance remained within the optical window (700 - 900 nm) where endogenous light absorbance of biological tissues is lower, making them good candidates for cell labelling (19).

### 3.2 Gold labelling of human umbilical cord mesenchymal stromal cells

Next, we assessed the effect of different GNR concentrations on morphology, labelling efficiency and cell viability. No overt changes in cell morphology were observed via microscopy 24 h after GNR labelling (Figure 2a, top). Using the gold-specific silver staining, GNR uptake by the hUC-MSCs was confirmed at all concentrations, showing a clear dose-dependent uptake trend as indicated by an increase in contrast. The gold particles accumulated in the perinuclear space, consistent with lysosomal accumulation as previously reported (Figure 2a, bottom) (20, 21).

**Figure 2.**
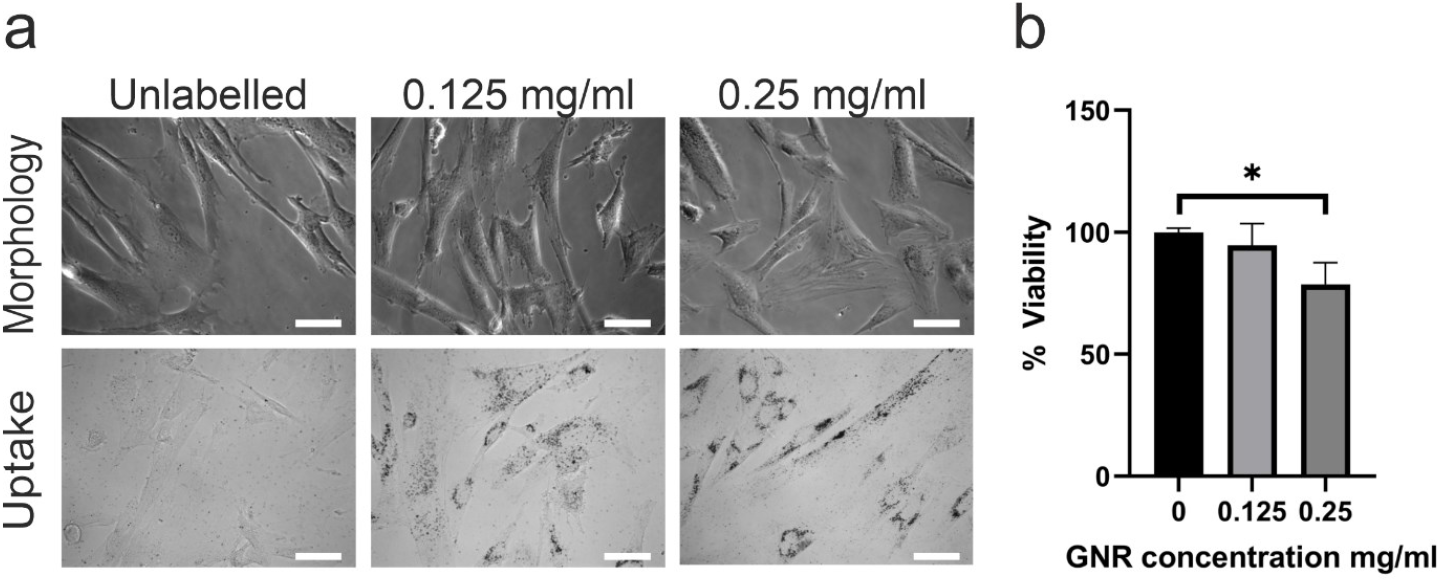
Cell morphology, viability and nanoparticle uptake after GNR labelling. (a) Cell morphology (top) after labelling with GNRs at different concentrations and uptake (bottom) assessed by silver staining. Optical microscopy images of hUC-MSCs labelled with different GNR concentrations for 24 h. Dark contrast is generated by silver-enhanced staining of GNRs. Scale bar 100 μm. (b) Cell viability. unpaired T-test p<0.05. n=3

To determine cell viability, we quantified the total amount of ATP in cells labelled for 24 h with 0.125 mg/mL or 0.25 mg/mL GNRs. Our results indicated that viability levels were at 94.6 %, and 78.7 % of unlabelled control cells. While a significant reduction in viability was observed at the highest concentration, the GNRs were not overtly toxic to the cells (Figure 2b). Given that labelling with 0.25 mg/mL yielded more uptake (Figure 2a, bottom right), this concentration was taken forward for cell phantom imaging with MSOT and CT.

### 3.3 MSOT/CT imaging of GNR labelled hUC-MSCs in phantoms

Before in vivo imaging, it is important to establish whether the GNR-labelled MSCs can be visualized by MSOT and CT. To do this, 5 × 10^5^ hUC-MSCs were suspended in 100 μL PBS (GNR-labelled and control MSCs) into an agar phantom and MSOT intensity was recorded at wavelengths ranging from 680 to 980 nm. The absorbance spectrum measured with the MSOT instrument was broadened compared to the spectrum measured with a suspension of dispersed gold nanorods. This may indicate loss of the silica shell, aggregation after uptake, and plasmon coupling (Figure 3a, left) (13). Nevertheless, labelled cells could still be detected after applying a multispectral unmixing algorithm, where they are seen as a crescent shape due to the cells sedimenting to the bottom of the phantom (Figure 3a, right). Imaging of the phantom by CT showed no difference in contrast between unlabelled and GNR-labelled MSCs when using either the standard filter (aluminium) or a specialized copper filter for the detection of metals (Figure 3b).

**Figure 3.**
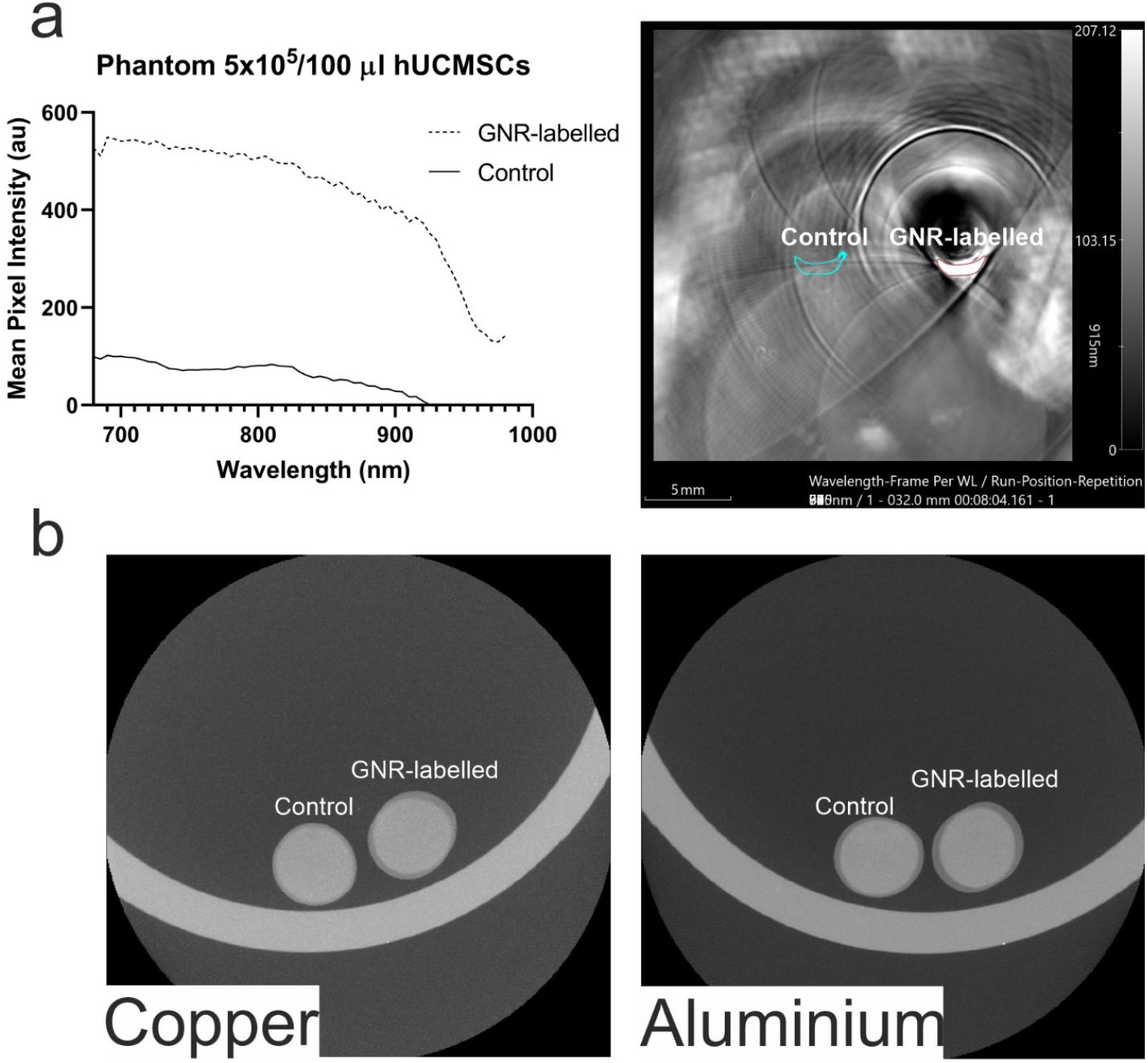
Phantom imaging of GNR-labelled and unlabelled hUC-MSCs. (A) MSOT demonstrates a clear distinction in signal intensity between samples containing labelled or unlabelled cells. The left panel shows the spectrum of GNR-labelled and unlabelled MSCs. The right panel shows a maximum intensity projection of imaging phantoms containing MSCs. (B) CT fails to detect GNR-labelled MSCs, with both samples having identical contrast regardless of the imaging filter used. Copper (left); Aluminium (right).

### 3.4 In vivo multi-modal monitoring of GNR-labelled cells administered subcutaneously or intravenously

To further investigate the potential for cell tracking of gold-labelled MSCs by MSOT and CT, 5 × 10^5^ control hUC-MSCs cells or hUC-MSCs labelled with 0.25 mg/mL GNRs were administered either IV or SC.

BLI demonstrated that following either delivery route, the hUC-MSCs showed a strong luminescent signal confirming their presence in the mice. SC injection resulted in the cells localising at the site of injection, with comparable signals at the site of unlabelled (left flank) or gold-labelled cells (right flank). By contrast, when the cells were administered IV, the hUC-MSCs localised to the lungs (Figure 4a).

**Figure 4.**
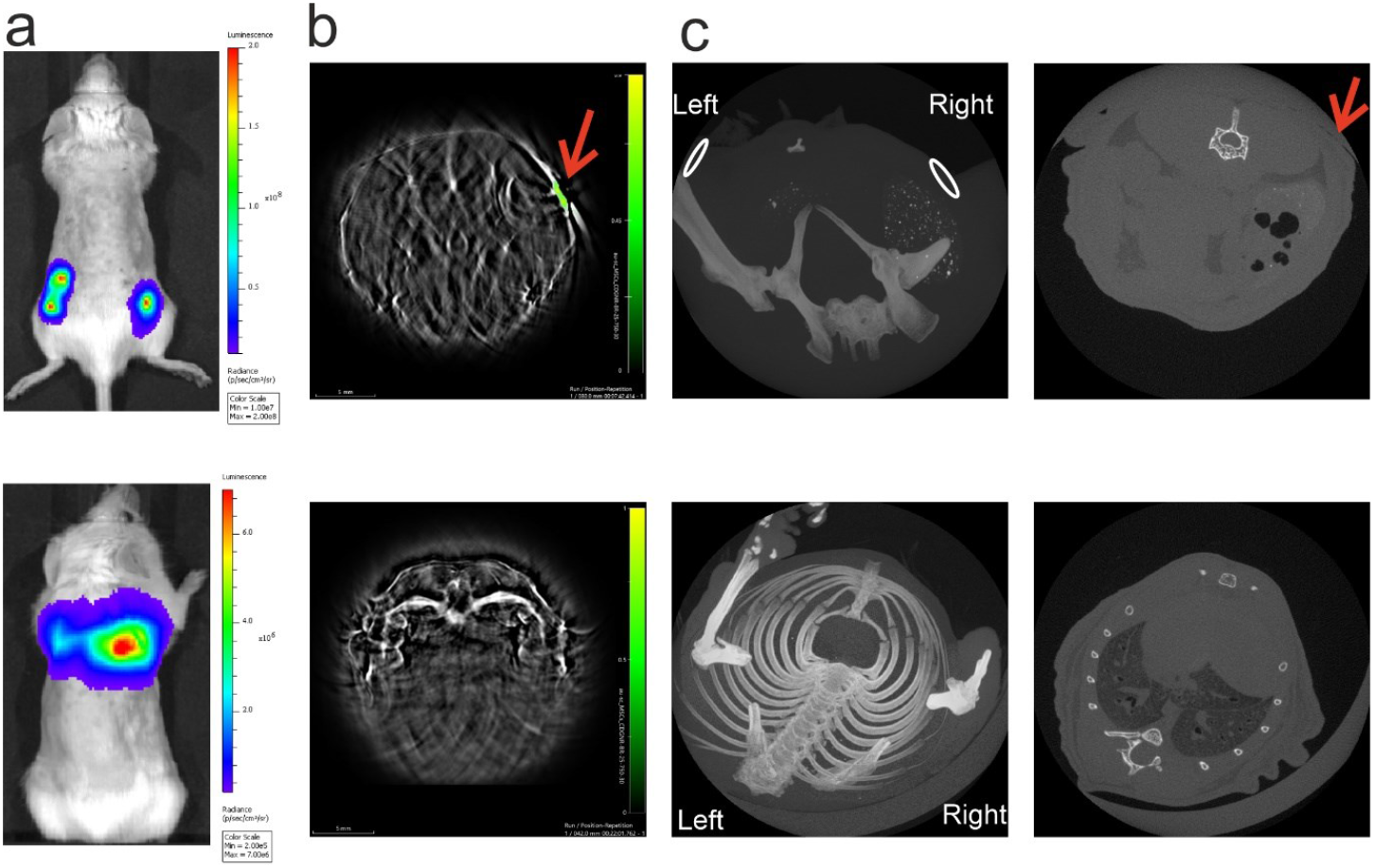
Representative bioluminescence, MSOT and CT imaging of mice after receiving hUC-MSCs. n=6 (a) BLI shows that cells injected SC remain at the site of injection whereas when injected IV, they localise to the lungs. (b) MSOT imaging of mouse after IV or SC injection. GNR labelled cells were distinguished from any internal organ when injected SC (green scale displays GNR-specific signal, indicated by arrow) but not IV. (c) CT fails to generate contrast of GNR labelled cells administered via either route. 3D volume rendered images (left), site of cell location is indicated by ovals. 2D representative CT section of the site of SC injection (top, right; indicated by a red arrow) and lungs (bottom, right).

MSOT confirmed the presence of GNR-labelled cells in the mouse right flank as observed by the high contrast resulting from the GNRs. On the other hand, the unlabelled cells failed to generate optoacoustic contrast, demonstrating the ability of MSOT to detect gold-labelled cells. As expected, the GNR-labelled cells in the lungs could not be detected due to the high air content within this organ (Figure 4b).

Finally, the SC injected GNR-labelled hUC-MSCs could not be detected by CT. A clear image of the lungs could be obtained by CT but the GNR-labelled MSCs were undetectable (Figure 4c).

These results demonstrate that 5 × 10^5^ luciferase-expressing hUC-MSCs can be detected by BLI following both IV and SC administration, gold-labelled hUC-MSCs can be detected by MSOT following SC administration, but CT lacks the sensitivity to detect the hUC-MSCs via either administration route.

## Discussion

One aim of multi-modal imaging strategies is the analysis of the whole-body and intra-organ biodistribution of administered cells in preclinical animal models. In this study, we explored the feasibility of combining the sensitivity of BLI with the spatial resolution of MSOT and the ability to image the lungs by CT to track GNR-labelled MSCs in vivo.

Photoacoustic imaging uses contrast that can be endogenous (for example due to absorption by haemoglobin) or exogenous via the use of contrast agents (13). For cell tracking by MSOT, labelling with gold nanorods has been a method of choice as these particles have a longitudinal plasmon band with strong absorption in the near-infrared (13,22–24). Therefore, the silica-coated GNRs used in this study, with LSPR bands at 738 nm or 713 nm in cell medium are good candidates for generating contrast in cells.

Previously, our group showed that a BLI/MSOT strategy to monitor intracardially administered GNR-labelled FLuc+ mouse MSCs revealed the presence of cells in the head, liver and kidneys of mice with both imaging techniques, proving the efficacy of this methodology for cell tracking (15).

Here, in line with earlier publications, BLI showed that IV injection leads to hUC-MSC accumulation and trapping in the lungs (1–3). Due to differences in sound propagation in air, the capacity of MSOT to detect GNR-labelled cells within the lung is hampered. However, following subcutaneous injection, GNR-labelled hUC-MSCs could be detected by MSOT.

CT has been used to track gold-labelled cells in the lung, as well as in other organs in vivo and ex vivo. Cell labelling strategies to achieve this vary widely (table 1). Studies describe using gold nanoparticles with different surface chemistries, diameters, initial gold concentrations, and incubation times (25–36). It has been shown that the properties of gold nanoparticles impact cell uptake (25), which agrees with the variation in uptake efficiency reported in these studies. Moreover, cell number, administration routes and injection volumes, tracking time, CT scanners, and scanning settings differ greatly between studies. This reflects the complexity of comparing results in the field of gold-labelled cell tracking by CT. Nonetheless, at least twelve reports suggest that CT enables longitudinal tracking of gold-labelled cells (25–36), although it should be noted that at least 6 of these originate from the same research group.

**Table 1.** Overview of articles reporting pre-clinical CT tracking of gold labelled cells. IPF = Idiopathic pulmonary fibrosis. RCS = Royal college of surgeons. *Gold nanoparticles (GNP) characteristics in order of appearance are size, coupling/coating, other characteristics. CPP= cell penetrating peptide. PSD= polymer polysulfonamide. PEG= polyethylene glycol, TAT= trans-activator of transcription. RBITC= [Rhodamine B isothiocyanate](https://www.sigmaaldrich.com/US/en/product/aldrich/283924). PLL= Poly-L-Lysine. DMSA= 2,3-dimercaptosuccinic acid. Y = yes. N = No.

| Cell type | Cell number | Animal model | Administration route | Tracking time | Labelling conditions | GNP characteristics* | Gold/cell | Imaging | Cells detected? | Ref |
| --- | --- | --- | --- | --- | --- | --- | --- | --- | --- | --- |
| hUCMSCs | 4 x 10<sup>6</sup> | Mouse IPF | Tracheal infusion | 1, 9, 25, 35 days | 100 µg/mL<br>4 h | 127.3 nm, CPP-PSD<br>pH-sensitive | 313.5 pg | *In vivo* | Y | (25) |
| hUCMSCs | 4 x 10<sup>6</sup> | Mouse IPF | Tracheal infusion | 1, 4, 7, 10 days | 1000 µg/mL<br>≥ 12 h | 40 nm, PEG-TAT<br>RBITC-labelled | 920 pg | *In vivo* | Y | (26) |
| hUCMSCs | 4 x 10<sup>6</sup> | Mouse IPF | Tracheal infusion | 1,3,5,7,10days | 200 µg/mL<br>12 hours | 40 nm,<br>temperature responsive | 120 pg | *In vivo* | Y | (27) |
| hUCMSCs | 1.5 x 10<sup>6</sup> | Mouse IPF | Tracheal infusion | 3,48 h, 9, 16, 23 days | 200 µg/mL<br>24 h | 10.7 ± 1.7 nm, BSA-PLL | 293 pg | *In vivo* | Y | (28) |
| Rat glioma C6 cell line | 1 x 10<sup>5</sup> | Wistar rat | Stereotactic injection into brain | 16 days<br>Ex vivo | 50 µg/mL<br>22 hours | 50 nm,<br>colloidal | 0.04 ng | *Ex vivo* | Y | (29) |
| hUCMSCs | 2 × 10<sup>5</sup> | RCS rat model | Subretinal injection | 1, 15, 30 days post injection | 1.4 × 10<sup>8</sup> particles/mL<br>24 h | 80 nm,<br>colloidal | Not mentioned | *In vivo* | Y | (30) |
| Human periodontal ligament stem cells (hPDLSCs) | 1 x 10<sup>6</sup> | Wistar rats | - Intramuscular<br>- Subcutaneous<br>- Submucosal<br>- Subgingival | 0, 2, 5 days | 50 µg/mL<br>12 h | 40nm, PLL-hydrobromide<br>RBITC-labelled | Not mentioned | *In vivo* | Y | (31) |
| hMSCs (undetermined source) | 2 × 10<sup>4</sup> or 5 × 10<sup>5</sup> | Sprague Dawley rats | Stereotactic injection into brain | 30 min post injection | 100 µg/mL<br>12 h | 40 nm, PLL<br>RITC-labelled | 382.5 pg | *In vivo* | Y | (32) |
| Bone marrow MSCs | 1 x 10<sup>6</sup> | BL/6 IPF | Tracheal infusion | 7, 14, 21 days | 200 µg/mL<br>24 h | 12.2 nm ± 1.59 nm, Albumin-PLL<br>ICG-labelled | 218 pg | *In vivo* | Y | (33) |
| hMSCs (undetermined source) | 1 x 10<sup>6</sup> | Nude mice | Intra-arterial (carotid artery) | 24 h | 52 µg/mL<br>22 h | 50 nm,<br>colloidal | 332 ± 2 µg | *Post mortem* | Y | (34) |
| F98 rat glioma cells | 1 x 10<sup>5</sup> or 2 x 10<sup>5</sup> | Nu/nu mice | Stereotactic injection into brain | 6-8 days after tumor implantation | No dose mentioned<br>4 h. | 50 nm,<br>Colloidal | 26,500 GNPs | *In vivo* | Y | (35) |
| Dental pulp MSCs | 5 x 10<sup>5</sup> and 1 × 10<sup>6</sup> | Mouse Silicosis | Intranasal | Daily up to 7 days | 90 µg/mL<br>24 h | 26.4 ± 0.96 nm, DMSA | 4 pg | *In vivo* | N | (36) |

In contrast, despite BLI and MSOT confirming the presence of cells in the mice, our study failed to detect contrast generated by the gold-labelled MSCs by CT regardless of whether they were delivered SC or IV. Silva *et al*. also reported failure to detect labelled cells in vivo by CT (36), and of the studies reported in table 1, this is the only one with negative results. They labelled hMSCs with dimercaptosuccinic acid (DMSA) gold nanoparticles (GNPs) for 24 h at a concentration of 90 µg/mL and observed a slightly higher contrast than unlabelled MSCs in CT phantoms, but the difference was not significant, and intranasal inoculation of labelled MSCs did not result in detectable contrast by in vivo CT.

Here, although not highly toxic, the gold concentration used showed reduced cell viability after 24 h. The fact that other groups have used higher labelling concentrations might be explained as GNP toxicity depends on functionalisation and uptake (37). The effects of surface modifications have been studied in various cell lines. Naked GNRs negatively affect mammalian cells at concentrations as low as 0.7 µg/mL while silica coating increases the cell tolerance to GNRs (38) consistent with the viability of hUC-MSCs exposed to the silica-coated GNRs in this study.

Silica shell thickness plays a key role in preventing plasmon coupling and preserving the GNRs optical signature after cell uptake, which are important considerations for optimal MSOT imaging (13). While the commercial GNRs used here lost their silica coating during labelling (contrarily to those used in (13)), MSOT still enabled the detection of the GNR-labelled hUC-MSCs in the mouse flanks. In contrast to MSOT, aggregation might work in favour of the detection of gold by CT by preventing the GNRs from being removed by exocytosis which would reduce intracellular gold (27,39,40). Despite this aggregation phenomenon potentially taking place in our study, the GNR hUC-MSCs were still undetectable by CT.

High gold uptake per cell is necessary to achieve good contrast as CT signal increases proportionally with increasing gold concentrations; however, cell uptake usually reaches saturation (41,42). The GNRs used here had an average core size of 56 × 18 nm, which results in high uptake by receptor-mediated internalization (43). We used the highest concentration that did not induce overt toxicity. Despite this, no CT signal was detected in our study.

The discrepancy between our results and those of other studies raises the question of the limit of detection of CT for gold. This determination is not straightforward as CT scanning conditions along with the properties of the gold nanoparticles impact X-ray attenuation (44).

Attempts at determining the minimal amount of gold necessary to achieve contrast in CT have been made using phantom imaging. Galper *et al*. established that the attenuation of gold is 5.1 HU/mM (45). This is a physical parameter that should not vary between research groups. We attempted to evaluate the attenuation of gold corresponding to results in the publications reporting cell tracking with CT. Table 2 shows those estimated HU/mM attenuations. To arrive at these numbers, we calculated the molar concentration of gold in the CT phantoms used in the studies using equation 1:

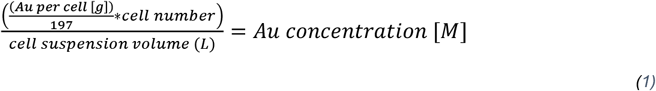

It is noteworthy that most studies show a much higher HU/mM attenuation in their phantom studies when compared to Galper’s data. Considering the 5.1 HU/mM attenuation, Cormode and colleagues concluded that 5.8 mM is the minimum detectable gold concentration (46). Considering a cell volume of 8 pL, the minimum amount of gold per cell necessary to achieve a 5.8 mM (1.16 g/L) concentration for a voxel filled entirely with labelled cells is 9 pg of gold/cell. Equation 2:

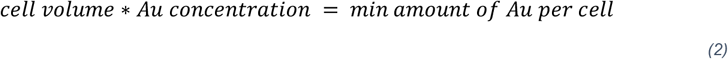

The gold/cell column in table 2 shows that all studies except for Silva *et al*. (36) achieved a nanoparticle load per cell higher than the 9 pg detection threshold, potentially explaining why Silva’s study is the only one that failed to visualize the gold-labelled cells by CT. It is thus clear that extremely high cellular uptake of GNPs is necessary in order to obtain CT contrast, which increases costs and may adversely affect cell health.

**Table 2.**
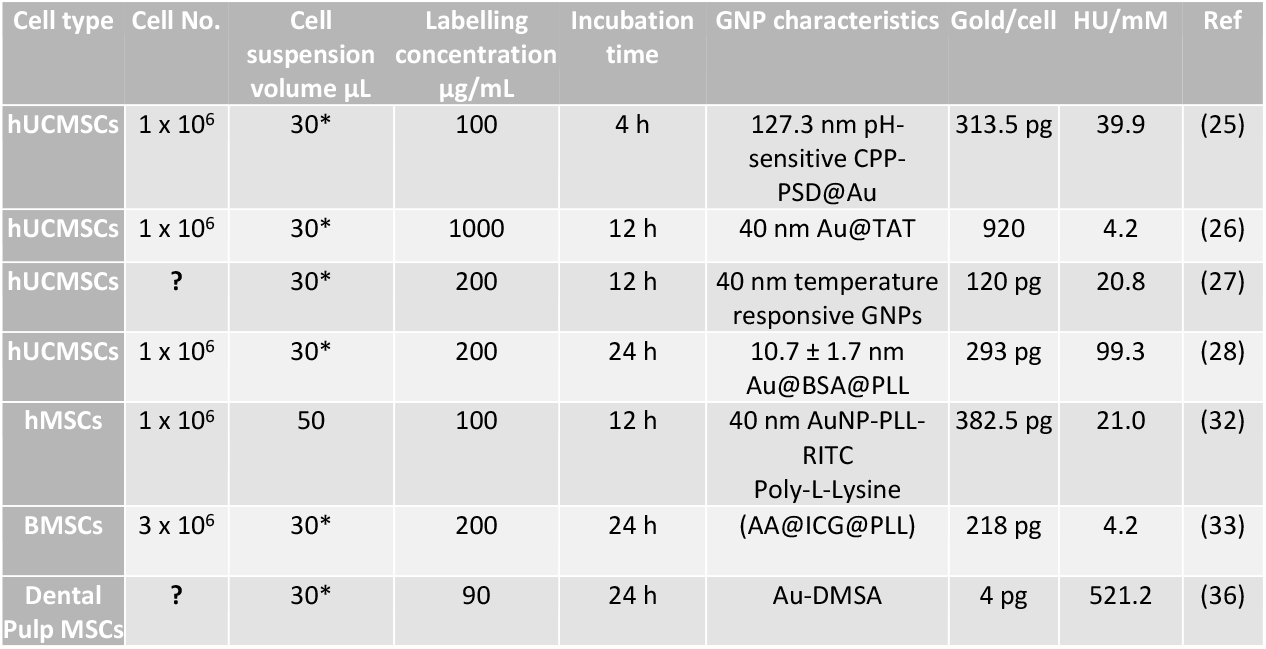
HU/Mm attenuation estimated from the published literature where CT imaging of gold-labelled cell phantoms was undertaken. Question mark (**?**) indicates that the cell No. was assumed to be 1 × 10^6^ cells as this information was not indicated in the papers. Aterisk (*) indicates that the cell suspension volume was estimated based on a pellet of 1 × 10^6^ cells.

The main limitation of our study is that we did not quantify the amount of gold per cell. Given the lack of contrast observed during the imaging of cell phantoms as well as in vivo, it is clear that even at the highest labelling concentration, the GNRs did not accumulate in high enough numbers inside the cells and thus, were not detectable by CT. On the contrary, they were easily detectable in the same conditions by MSOT when injected subcutaneously, showing that this imaging modality is significantly more sensitive than CT.

## 4 Conclusion

We tested the feasibility of a non-invasive, multimodal imaging approach that utilises a combination of GNRs and reporter genes to track MSCs after subcutaneous or intravenous injection in vivo. This labelling approach did not affect cell morphology and viability of hUC-MSCs significantly and allowed for robust tracking of the cells by BLI for both IV and SC delivery. The GNR-labelled cells were detectable by MSOT when injected subcutaneously validating the ability of a BLI/MSOT tracking approach. Although CT produces anatomical images of the lungs, the same GNR-labelled cells could not be detected within this organ or in the flanks of the mice indicating that the cells did not carry enough contrast agent to be tracked by CT.

To provide enough contrast for CT imaging, large amounts of gold are necessary. However, high labelling concentrations might impair cell viability making CT tracking of gold labelled cells challenging.

In summary, our study found that multimodal imaging of MSCs labelled with gold nanoparticles and the reporter gene firefly luciferase allows BLI and MSOT detection of administered cells in vivo, however CT lacks sensitivity toward gold under the conditions investigated.

## Data availability

*All datasets from this study are publicly available on Zenodo, http://doi.org/10.5281/zenodo.6624805*

## Conflicts of interest

James Littlewood works at iThera Medical GmbH. The other authors declare that they have no conflicts of interest in relation to this work.

## Acknowledgements and funding statement

In vivo imaging data in this article were obtained in the Centre for Preclinical Imaging (CPI) of the University of Liverpool. The CPI has been funded by a Medical Research Council (MRC) grant (MR/L012707/1). We thank Ms Alison Beckett (Biomedical EM Unit, University of Liverpool) for assistance with electron microscopy imaging, Dr Tim Devling and Dr Neal Burton (iThera Medical) for expert advice on MSOT imaging. MSC transduction and sorting was carried out by Mr Francesco Amadeo, NHSBT. This project has received funding from the European Union’s Horizon 2020 research and innovation programme under the Marie Skłodowska-Curie grant agreement No. 813839 and from the European Research Council (ERC) under the European Union’s Horizon 2020 research and innovation programme (grant agreement No. [856478]).

## Notes

http://doi.org/10.5281/zenodo.6624805

